# Complementary roles among traits in community assembly processes of benthic foraminifera

**DOI:** 10.64898/2026.11.19.767260

**Authors:** Galli Giacomo, Jonkers Lukas, Woodhouse Adam, Hernández-Almeida Ivan, Diz Paula, Schweizer Magali, Morigi Caterina, Sabbatini Anna, Cannings Torin, Gamboa-Sojo Viviana, Mojtahid Meryem, Nardelli Maria Pia, Swain Anshuman

**Affiliations:** Department of Environmental Science, Informatics and Statistics, University of Venice; Department of Earth Sciences, University of Pisa; Max Planck Institute for Chemistry, Department of Climate Geochemistry, Mainz, Germany; MARUM - Centre for Marine Environmental Sciences, University of Bremen, Bremen, Germany; School of Environment, Earth and Ecosystem Sciences, Open University, Walton Hall, Kents Hill, Milton Keynes, MK67AA, U.K; Past Global Changes International Project Office – University of Bern, Hochschulstrasse 4, Bern, 3012, Switzerland; Centro de Investigación Mariña, XM1, Universidade de Vigo, Vigo, Spain; Univ Angers, Nantes Université, Le Mans Université, CNRS, Laboratoire de Planétologie et Géosciences, LPG UMR 6112, 49000 Angers, France; Department of Life and Environmental Sciences, Marche Polytechnic University, Ancona, Italy; Section of Earth and Environmental Sciences, University of Geneva, Switzerland; Central American School of Geology, University of Costa Rica; Department of Ecology and Evolutionary Biology, University of Michigan, Ann Arbor, Michigan 48109, USA; Museum of Paleontology, University of Michigan, Ann Arbor, Michigan 48109, USA

**Author notes:** **Authorship:** GG wrote the main text and performed the analysis; all authors have substantially contributed to the revisions and development of the work.

**Keywords:** Community ecology, Benthic foraminifera, benthos, network, morphological traits

## Abstract

Organismal traits play a distinct role in shaping community assembly, yet they have received comparatively little attention in marine communities. Here, we focus on living benthic foraminifera, a diverse group of marine protists that inhabit the seafloor. Benthic foraminifera are protists that have evolved different test-construction strategies among which select and agglutinate grains from their surrounding environment or by biomineralize. In addition, taxa exhibit distinct morphologies have been hypothesized to reflect microhabitat partitioning within sediments. Here, we investigate their metacommunity patterns by analysing a co-occurrence network covering the Atlantic and Pacific Oceans. We found that test-construction strategies significantly explained the structure of the network, whereas the general test shape did not. We hypothesize that test-construction strategy act as a broader-scale filter on the regional species pool, whereas morphological differences may be more closely related to local-scale coexistence and habitat partitioning. This study supports the idea that different organismal traits have different roles in the community assembly process.

## 1 Introduction

How ecological communities assemble across broad spatial and temporal scales remains a fundamental yet elusive question in ecology and evolution (Vellend, 2010). Although much of our understanding of metacommunity processes has developed from theoretical frameworks and modelling approaches, our empirical knowledge comes only from a few groups of mainly terrestrial organisms (Logue *et al*. 2011). Consequently, there is a need for empirical studies that integrate organismal traits with metacommunity processes across broad spatial scales and environments to generalize our understanding of how ecological communities assemble (Leibold *et al*. 2004; Logue *et al*. 2011). Morphological traits are key attributes by which organisms interact with their environments, and trait-based approaches provide insights into the ecological mechanisms underlying community assembly (McGill *et al*. 2006). However, how exactly organismal characteristics translate into patterns of community structure and biodiversity across spatial and temporal scales remains incompletely understood across diverse taxonomic groups and environmental realms (Mayfield & Levine 2010; Spasojevic & Suding 2012). In addition, traits can be challenging to define and quantify consistently, particularly when morphological characteristics do not directly capture the ecological functions or environmental affinities of organisms. Shelled organisms may be particularly well suited to investigating the relationship between morphological traits and community assembly, as their traits can be directly characterized from shell architecture. Thus, by focusing on marine shelled communities, we can address a fundamental question in community assembly theory: do different traits have different roles in the assembly and structure of ecological communities?

Benthic foraminifera (BF, phylum Foraminifera, (Adl *et al*. 2019)) are ubiquitous marine protists, with some freshwater and soil representative that potentially provide a simple biological system to ask how different organismal traits influence the rules governing community composition across different scales, and, more importantly, whether different traits play different roles in determining their spatial patterns. Dispersal is highly efficient in this group of organisms, promoting the transport of the resting juvenile forms (propagules) across a plethora of environments (Alve 1999; Buzas *et al*. 2015; Goetz *et al*. 2022; Pawlowski *et al*. 2007). While every taxon could be virtually anywhere, identical propagule banks gave rise to different community compositions depending on different environmental conditions (Buzas-Stephens *et al*. 2025; Goldstein & Alve 2011). Moreover, understanding how BF communities’ assembly has implications beyond their biology and ecology, since they play fundamental roles in marine benthic ecosystems, regulating the transfer of energy between different trophic levels as well as being a fundamental component in the carbon cycle (Buzas-Stephens *et al*. 2025; Gooday 1993; Moodley *et al*. 2000; Nomaki *et al*. 2008; Włodarska-Kowalczuk *et al*. 2013). Understanding what drives their spatial patterns is thus fundamental to properly understands associated changes in marine benthic ecosystem.

Within BF, functional traits could be directly related to the test morphology (Guy-Haim & Bouchet 2025). The fact that their test can be preserved in the geological record has made foraminiferal communities exceptionally useful across a wide range of disciplines, from modern ecology to paleontology (Gooday 2003; Murray 2006). Some taxa biomineralize calcium carbonate to produce calcareous tests of different mineral composition, whereas others select and agglutinate sedimentary grains from the environment to form agglutinated tests (Murray 2006). Additionally, different lineages have converged on similar solutions, either by biomineralizing calcite through metabolic pathways (Fouet *et al*. 2024; Henry & Basso 2026; Hoober *et al*. 2026; Pawlowski *et al*. 2003, 2013) or by using their reticulopodia to select and agglutinate different types of sediment grains (Murray *et al*. 2011). Besides test construction mechanisms, the general shape of the test has been recognised as another important trait associated with different ecological strategies (Corliss 1985; Gooday & Jorissen 2012; Guy-Haim & Bouchet 2025; Jorissen *et al*. 1995; Schmiedl *et al*. 2023). Test shape, broadly classified into ovoid, rounded, irregular, and elongated forms, has been associated with microhabitat partitioning. Although exceptions occur, elongated forms are generally associated with infaunal lifestyles (within seafloor sediments), whereas more rounded forms tend to be associated with epifaunal lifestyles (at the sediment-water interface, on hard-substrates or algae) (Schweizer *et al*. 2011; Van Der Zwaan *et al*. 1999) . By analysing co-occurrence patterns using a network approach, this study proposes that test-building strategies and general test shape have different roles in structuring living BF metacommunities, suggesting that different traits can contribute differently to community assembly processes.

## 2 Methods

To investigate the macroecological patterns in BF, an integrated statistical framework linking genus-level trait information, community composition, and network-based inference of trait associations was used. All the statistical and network analyses were performed in the R environment (R Core Team, 2024).

### 2.1 Dataset

We integrated the BENFEP_v2 (Diz *et al*. 2023) with the dataset of Murray (2006), both of which have been used extensively in other studies and publications (Buzas *et al*. 2023; Diz *et al*. 2024; Jones & Murray 2017). The dataset comprises 628 sites and 297 genera covering the Atlantic and Pacific oceans, between 73.75°N and 56.32°S, within a gradient of water depth between 0.3 and 6250 m (Fig. S1). Detailed information about the dataset is given in the supplementary material. Due to the differences in operators retrieving the data (Diz *et al*. 2023; Murray 2006) and since BF have been systematically defined by morphogroup approaches (Loeblich & Tappan 1988; Murray 2006), we decided to focus on the genus-level rather than species-level communities. This approach is justified given that taxonomic redundancy at the species level may arise when multiple taxa provide overlapping ecological information or when taxonomic distinctions are uncertain (Roberts *et al*. 2016). In BF, higher taxonomic levels retain much of the signal observed at the species level and can be easily transposed into paleontological studies (Balmford *et al*. 1996; Bevilacqua *et al*. 2012; Liu *et al*. 2023; Mistri & Rossi 2001). Moreover, our analysis focused mainly on rather stable characteristics of the organisms, namely how the test is formed and its general shape (Loeblich & Tappan 1988; Murray 2006) . Thus, by focusing on the genus level, we removed a negligible fraction of functional diversity, while retaining the most stable features of these communities. Due to the scarcity of data on BF taxa with organic-walled tests (e.g., allogromiids) we did not include them in the analysis, even though they represent important constituents of BF communities in deep-sea settings (Li *et al*. 2025; Nguyen *et al*. 2026; Pawlowski *et al*. 2013; Sabbatini *et al*. 2025; Song *et al*. 2026).

Traits were attributed at the genus level following the holotype description and by consulting standard BF taxonomic atlases (Burkett *et al*. 2024; Holbourn *et al*. 2013; Loeblich & Tappan 1988). The list of all the taxa and the assigned traits (test-construction strategies and test-morphology) is given in the supplementary materials.

### 2.2 Statistical analysis

To understand the roles that different morphological traits might have in shaping the overall metacommunity patterns in BF, we focused on the co-occurrence patterns of different taxa by analysing the network structure. Co-occurrence networks (both positive and negative) can provide a powerful framework to understand different community assembly mechanisms because patterns of taxa associations can be related to shared environmental preference, niche partitioning processes, ecological interactions and/or eco-evolutionary processes, all of which translates into the network topology (Tylianakis & Morris 2017).

Before building the network, we analysed the datasets for potential spatial autocorrelations (Fig. S1). First, we calculated Moran’s eigenvector Maps (MeMs) of our sites (from the latitude and longitude) to better constrain the differences in spatial patterns that were present in the dataset (function *dbmem*, Dray 2016; Dray et al. 2006; Guénard & Legendre 2024). To remove redundancy in MeMs, a forward selection algorithm (function *forward.sel*, Dray, 2016) computed on the 25 total MeMs was used to retain only 14 of them (Figure S4). Afterwards, a Mantel test (function *mantel*, Oksanen et al. 2024) evaluated how community dissimilarity correlates with geography; while a second test examined its relationship with depth. In addition, we computed a variation partitioning analysis (function *varpart*, Oksanen et al., 2024) to assess the relative contribution of these two parameters. Next, we conducted a constrained Redundancy Analysis (pRDA, function *rda*) using depth and the 14 MeMs on the Hellinger-transformed abundances of the taxa (Oksanen et al., 2024). Prior to the analysis, taxa with maximum abundance that was not exceeding 5% were removed (93/297).

To consider the possible effects of geography and bathymetry when studying BF associations, residuals extracted from the pRDA were used to compute both positive and negative co-occurrence networks using Spearman correlations. A conservative threshold (Spearman’s |r| > 0.5 and Bonferroni-corrected p-value < 0.05), was used to draw undirected links between two taxa. Then, we computed modularity (Q, Louvain algorithm, Csárdi et al. 2006), to assess how clustered the networks were, and used a null model to assess its significant departure from randomness by simulating degree-conserving networks (n=999). Afterwards, three different Exponential Random Graph Models (ERGMs) were used to analyse the topological features of the networks (function *ergm*, Krivitsky et al. 2023; Silk & Fisher 2017). ERGMs were evaluated both for their differences in explanatory power (Akaike’s Information Criterion AIC, Bayesian Information Criterion BIC), along with a pairwise Analysis of Variance between different models (function *anova*, package vegan, Oksanen et al., 2024). In ERGMs, the edge term provides the baseline propensity for an edge to occur between two nodes before incorporating node attributes or other structural terms. Hence, a significant negative edge coefficient indicates that the probability of observing an edge is lower than expected under the corresponding baseline network structure. Attribute terms can then be interpreted relative to this baseline. When the coefficient for an attribute is positive and strongly significant (likelihood θ > 0, p < 0.05), nodes belonging to the same category have a greater propensity to form an edge than expected from the baseline, indicating homophily. Conversely, when the coefficient is negative and strongly significant (θ < 0, p < 0.05), nodes belonging to the same category have a lower propensity to form an edge than expected from the baseline. When the coefficient is not significant (p > 0.05), the attribute does not provide evidence that the propensity for an edge differs from the baseline. In both negative and positive networks, we fitted one model which considered only the test-formation property, one where only the morphology was considered and one where both effects were considered. Afterwards, we analysed the ability of the model to reproduce the network statistics by using goodness-of-fit analysis (function *gof*, Krivitsky et al., 2023).

## 3 Results

Both geographic distance and water depth were significantly associated with community composition (Mantel test, rMeM= 0.275, r_depth_= 0.255, both p < 0.001). Similarly, variation partitioning showed that geography alone explained 19% of the total variation, bathymetry alone explained 1%, and their shared contribution accounted for 6%, leaving 74% of the variability unexplained (Fig. 1a). To reduce the influence of these spatial and bathymetric effects, we performed a partial redundancy analysis (pRDA) constrained by MeMs and water depth (Fig. S5). Like the previous results, geography and bathymetry together accounted for 28% of the variation, whereas 72% of the variance remained unexplained in the pRDA (Fig. 1b). We then analysed the co-occurrence network derived from pairwise Spearman correlations calculated from the pRDA residuals (Fig. 1c). The positive network yielded 150 nodes, 594 edges and a significant high modularity (Fig. S7a, Q = 0.76, z = 63.46, p < 0.001), indicating a strong tendency for taxa to form distinct subcommunities (Fig. 1c). On the other hand, the negative network yielded 53 nodes, 135 edges and the modularity was not significant (Fig. S7b, Q = 0.29, z = -1.38, p = 0.911), providing no evidence of non-random modular structures, suggesting that negative associations were randomly compartmentalized. Since network modularity did not differ significantly from random expectations (Fig. S7), we based our subsequent inferences exclusively on the positive network. Furthermore, we assess whether the positive network topology was primarily related to taxon abundance and occupancy by examining the relationship between the mean relative abundance and occupancy (as defined as the proportion of sites occupied by a taxon) against the node’s degree and the participation coefficient (*sensu* Guimerà & Amaral 2005, Fig. S8). Mean relative abundance and occupancy were not significantly associated with the participation coefficient (Fig. S8a, c; Pearson’s r = -0.09, p = 0.27; r = -0.11, p = 0.2, respectively), but they were negatively associated with degree (Fig. S8b, d; Pearson’s r=−0.321, p<0.001; r= -0.28, p < 0.001, respectively). Together with the construction of the network from pRDA residuals, these results suggest that the observed modular organization is not simply a consequence of the major spatial and bathymetric gradients, taxon abundance, or occupancy. Instead, they support the idea that the network structure may reflect the influence of distinct ecological processes shaping community organization (Ponisio *et al*. 2019) .

**Figure 1.**
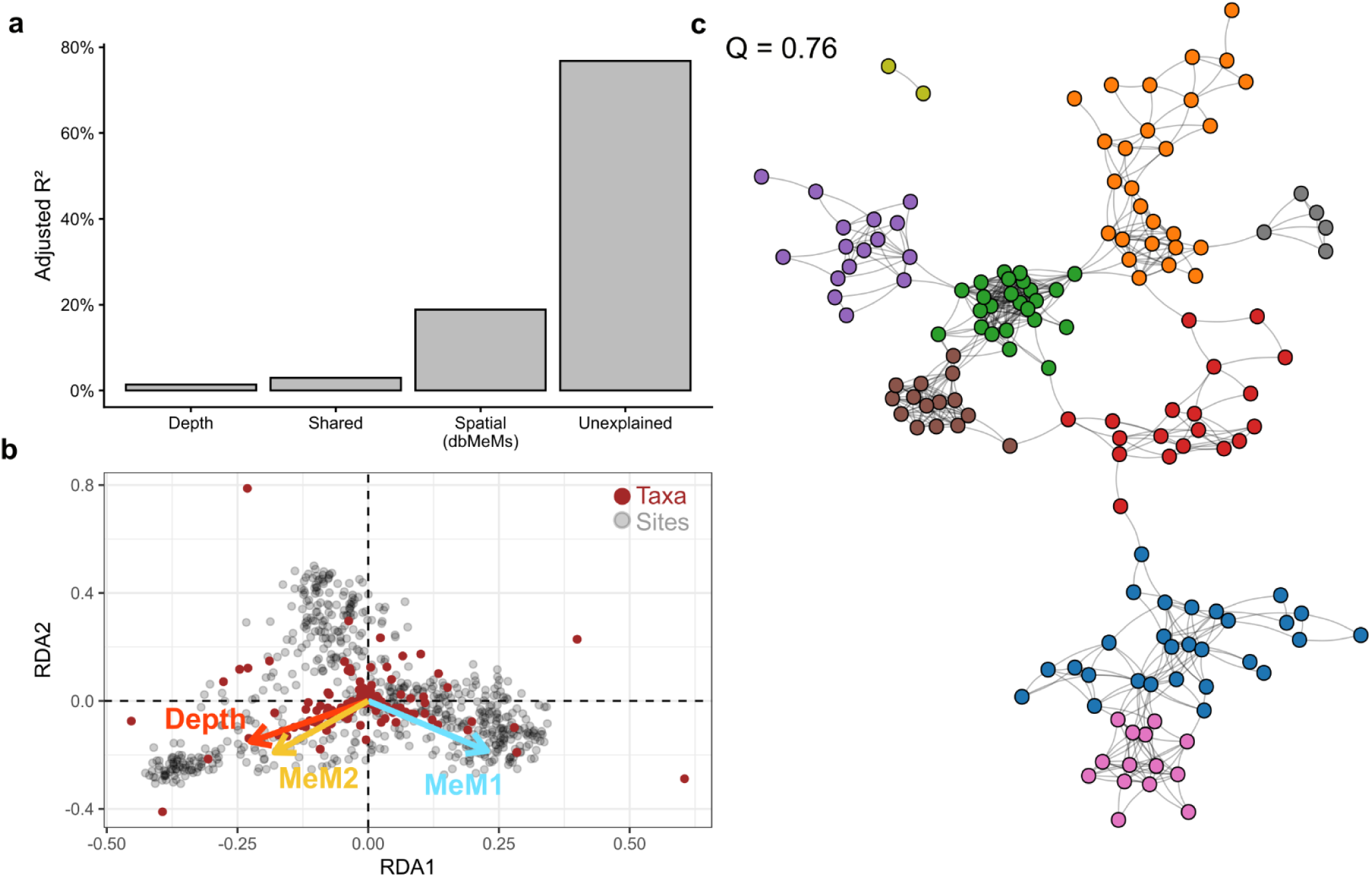
Metacommunity structure of living benhic foraminiferal community. a) Variation Partition results showing the variance explained by the bathymetry (depth), geography (dbMeMs) and the shared effect of both, as well as the unexplained variance. b) Biplot derived from the pRDA result. Only the depth (red arrow), MeM1 (light blue arrow) and MeM2 (orange arrow) (the other MeM are shown in Figure S5). c) Positive co-occurrence plots showing the associations of taxa evaluated on the residuals of the pRDA (panel b). Different colors represent different modules detected using the Louvain algorithm. Q refers to the modularity of the network. The network was plotted using the Kamada-Kawai algorithm.

The Exponential Random Graph Models (ERGMs) showed different results (Fig. 2, Table 1). All edge terms (the baseline) were highly significant and negative (Table 1), indicating the low propensity for an edge to form between two nodes. Test-building strategy showed a significant positive association (θ = 0.67, p<0.001), whereas test morphology did not (θ = 0.11, p = 0.184). The combined models yielded comparable results and did not significantly improve model fit (Fig. 2, Table 1, ANOVA: Δdeviance = 1.5, df = 1, p = 0.2132).

**Figure 2.**
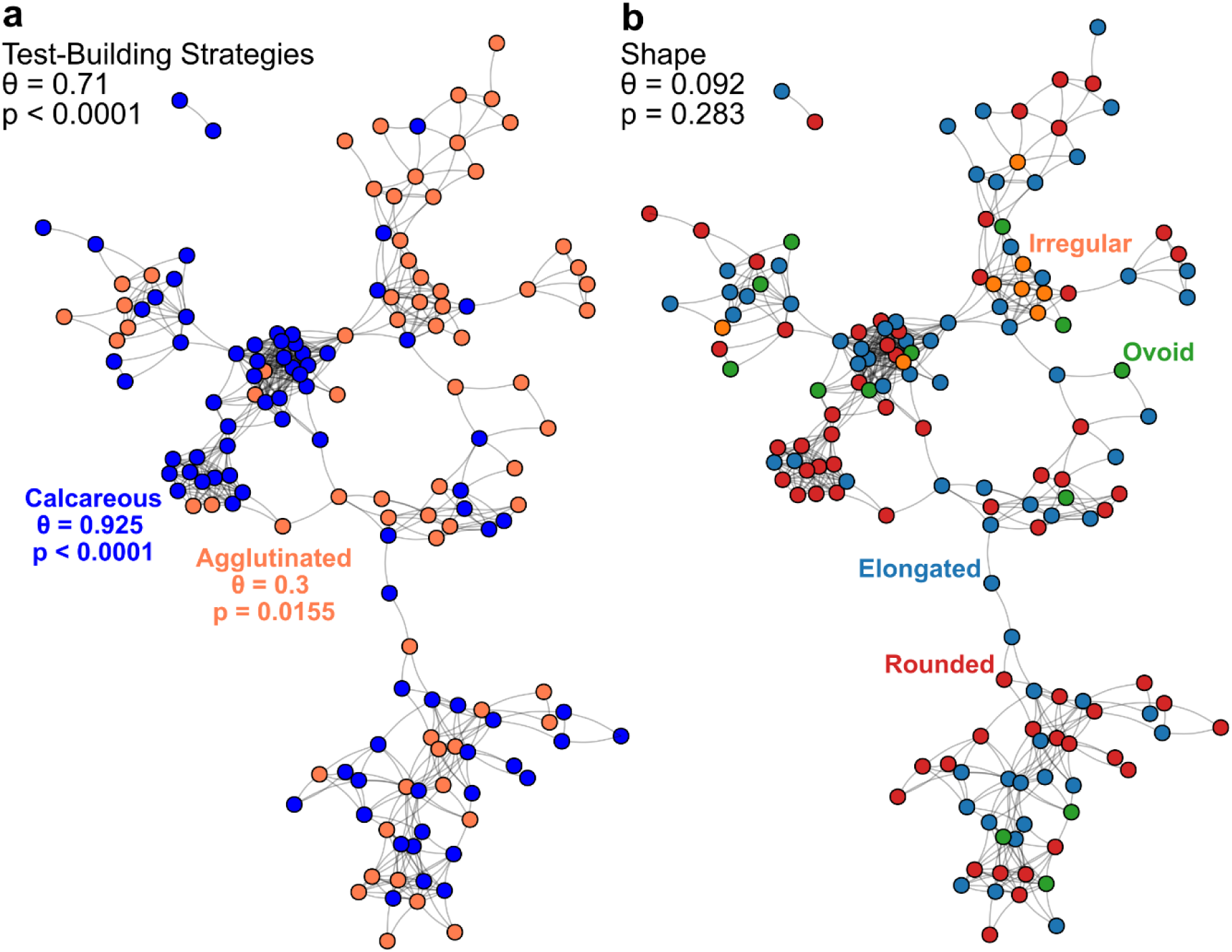
Traits distribution in the positive co-occurrence networks of benthic foraminiferal communities. a) Test-building strategies. Taxa showing calcareous tests are shown in blue, while agglutinated ones are shown in orange. b) General Shape of the test. Taxa with irregular-shaped tests are shown in orange, ovoid forms in green, elongated forms in blue and rounded forms in red. Likelihood θ and associated p-values of the ERGM models where both traits were considered (Further information in Table 1).

**Table 1.** Exponential Random Graph Models (ERGM) results obtained from the positive co-occurrence network and the negative co-occurrence network. The AIC refers to the Akaike’s Information Criterion calculated on the model, BIC stands for the Bayesian Information Criterion and θ refers to the log-likelihood of two nodes to form a tie (see Methods). In bold are highlighted the significant log-likelihood.

| **ERGM Model** | **AIC** | **BIC<sup>-</sup>** | **Terms** | **θ** | **Standard error** | **z-value** | **p-value** |
|---|---|---|---|---|---|---|---|
| **Test-building** | 4579 | 4593 | **Edge** | **-3.291** | **0.072** | **-45.707** | **<0.0001** |
|  |  |  | **Test-building** | **0.71** | **0.089** | **7.984** | **0.0019** |
| **Shape** | 4644 | 4658 | **Edge** | **-2.931** | **0.055** | **-53.658** | **0.0001** |
|  |  |  | Shape | 0.131 | 0.086 | 1.526 | 0.393 |
| **Test-building + Shape** | 4579 | 4601 | **Edge** | **-3.331** | **0.079** | **-42.050** | **<0.0001** |
|  |  |  | **Test-building** | **0.707** | **0.089** | **7.940** | **<0.0001** |
|  |  |  | Shape | 0.108 | 0.086 | 1.249 | 0.212 |

We therefore focused subsequent ERGM analyses on test-construction strategy. First, we evaluated if different test-construction strategies yield different likelihood when considered separately. By considering differences in the traits, the model fit significantly improved (AIC/BIC=4550/4572; ANOVA, Δdeviance = 31.3, df = 1, p < 0.0001). Both test-construction strategies showed homophily, but with fundamental differences: agglutinated forms showed a lower likelihood to form a tie (θ = 0.3, z = 2.47, p = 0.0155), while calcareous forms were characterized by a higher one (θ = 0.925, z = 9.74, p < 0.0001). However, goodness-of-fit analysis (Krivitsky *et al*. 2023) indicated that edge density and mixing patterns were reproduced, but not fundamental topological features of the observed network, such as the degree distribution, shared partners distributions and minimum geodesic distances (Fig. S9). We therefore evaluated the effect of structural terms as the geometrically weighted edgewise shared partners term (GWESP with varying decay parameters, 0.1, 0.5, 1), which refers to nodes sharing similar network partners (i.e., transitivity, Krivitsky et al., 2023). However, these models resulted in non-varying GWESP statistics and unstable Markov Chain Monte Carlo estimation, preventing inference from the structural terms. We interpret these discrepancies as evidence that different test-construction strategies do not fully explain the higher-order organization of the network structure.

## 4 Discussion

Metacommunity processes and network co-occurrence patterns are tightly connected since the latter is shaped by spatial, environmental and demographic processes (Leibold *et al*. 2004). Therefore, the contrasting organization of the positive and negative networks suggests that BF metacommunity structure may be primarily shaped by shared environmental requirements or habitat associations, rather than consistent antagonistic interactions, such as resource competition or competitive exclusions (Fig. 1c, (Blanchet *et al*. 2020; Grilli *et al*. 2016; Li *et al*. 2022a). This is consistent with local-scale studies that showed that environmental filtering, along with ecological drift, are the main drivers in shaping BF communities (Li *et al*. 2025; Nguyen *et al*. 2026). Our results thus extend this mechanism to the scale of ocean basins. Despite not fully capturing the high-order network structures (Fig. S9), ERGM results indicate that test-building strategy is more strongly associated with BF metacommunity organization than test morphology, providing evidence that different traits are related to different properties of the metacommunity organizations (Table 1). This supports the idea that, for BF, different traits may contribute to community assembly at different scales. Thus, we hypothesize that test-building strategy may contribute primarily to a broader-scale filter acting on the species pool, whereas test morphology may be more closely related to local coexistence and habitat partitioning (Fig. 3).

**Figure 3.**
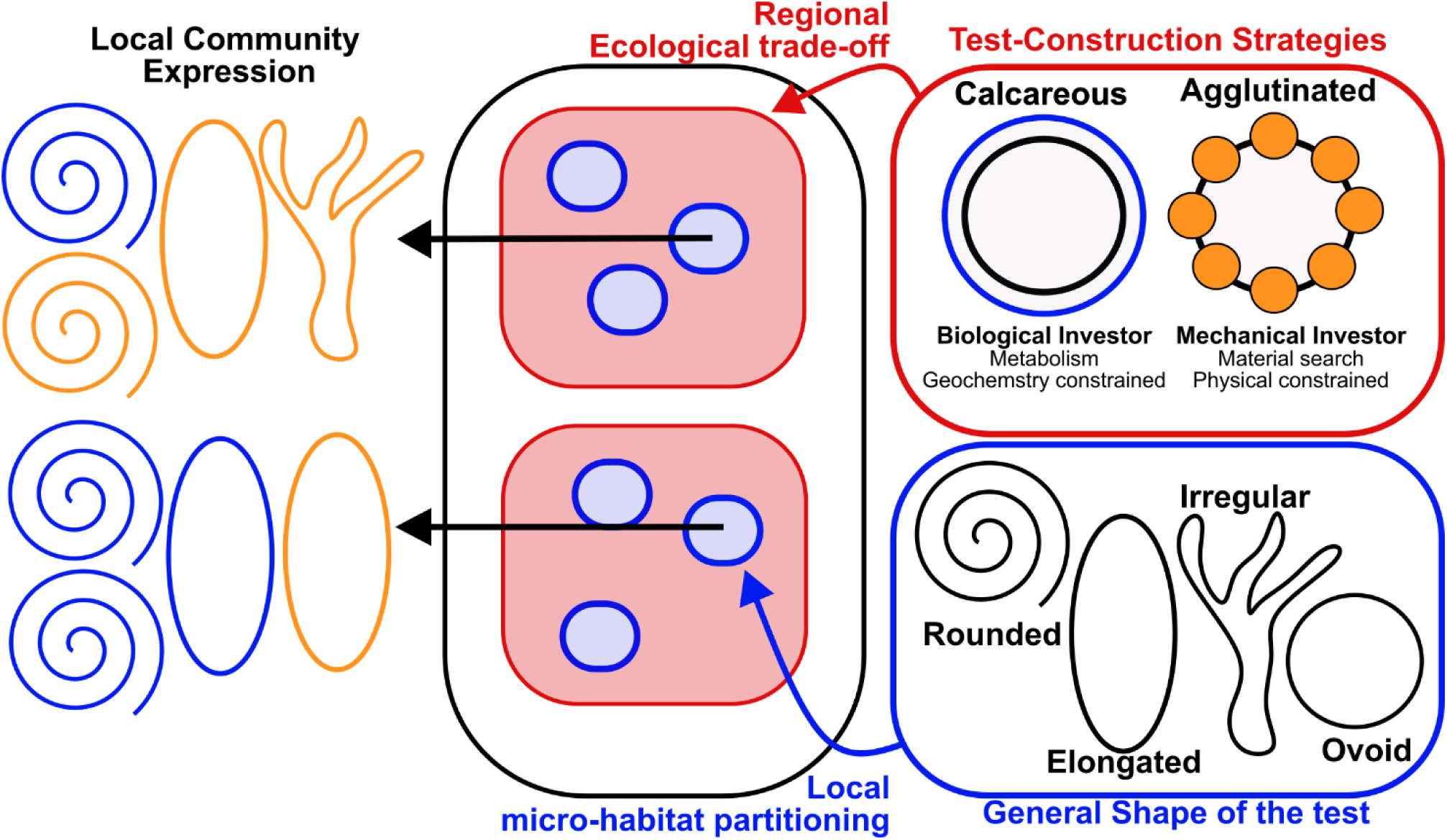
Hypothesized multi-level processes underlying benthic foraminiferal community assembly. For a benthic foraminiferal taxon to persist within a community, two ecological filters are proposed. A broader filter (red) imposes constraints on community structure through differences in energy allocation, metabolic pathways and constraints associated with test-building strategies (calcareous vs. agglutinated), represented here as mechanical investors versus biological investors. At the local scale, how the microhabitat is partitioned (blue), constrain test shape. The local community is thus expressed by the interplay between these two ecological filters.

Accordingly, differences between agglutinated and biomineralized construction strategies could influence the subset of taxa capable of occupying a regional environmental space. This process can be viewed conceptually through the Hutchinsonian fundamental niche (Holt 2009): physiological and metabolic differences associated with agglutination and biomineralization might constrain the environmental conditions under which taxa can potentially persist. Agglutinating BF can be thought of as mechanical investors (Fig. 3). They must identify, transport, and incorporate suitable particles into their tests, potentially requiring investment in processes associated with particle selection and manipulation (Bowser 2002; Gooday *et al*. 2007; Murray *et al*. 2011). Biomineralizing BF, in contrast, can be seen as biological investors (Fig. 3). They must allocate resources to the physiological and internal metabolic processes underlying mineral formation and maintenance (Courtial *et al*. 2025; Keating-Bitonti & Payne 2017; Prazeres *et al*. 2015). These different requirements may therefore impose different constraints on environmental persistence and community expressions, which is also consistent with the different homophilies detected within agglutinating and calcareous forms (Fig. 3).

We also infer that the general shape might operate more strongly at the local scale (Fig. 3). In BF, test morphology has been associated with different ecological strategies, namely microhabitat occupation, and interactions with the surrounding environment (Corliss 1985; Jorissen *et al*. 1995, 2022). Morphological differences may therefore influence how taxa partition the local environmental space in which they coexist. In this conceptual framework, the test-shape can be related to the realized niche, representing the subset of environmental conditions occupied after accounting for additional ecological constraints and interactions (Holt, 2009). Thus, we propose that test-building strategy and test-shape represent different components of a multi-level assembly process: construction strategy may constrain regional membership in the available pool, while test-shape may contribute to determining which taxa establish and coexist under local conditions (Fig. 3). Thus, different traits have different roles in BF community assembly: traits associated with fundamental physiological or constructional constraints may contribute primarily to regional species-pool formation, whereas traits associated with ecological strategy and microhabitat use may contribute more strongly to local community organization (Alve 1999; Buzas-Stephens *et al*. 2025; Li *et al*. 2025; Nguyen *et al*. 2026; Pawlowski *et al*. 2007).

Although this framework is developed here for benthic foraminifera, multi-level community assembly processes have been proposed across diverse taxonomic groups, including plants, microorganisms, fungi, fishes, and insects (de Bello *et al*. 2013; Davison *et al*. 2016; Lechêne *et al*. 2018; Luzuriaga *et al*. 2012; Neff *et al*. 2022; Saito *et al*. 2016). The recurrence of trait-mediated filtering across distantly related organisms and different environments further strengthens the possibility that these processes may represent a general feature of community assembly. Connections between community assembly processes and network structure have also been empirically documented in microbial communities and bipartite plant– insect interactions (Galai *et al*. 2024; Li *et al*. 2022b; Ponisio *et al*. 2019). Although previous studies have characterized both assembly processes and co-occurrence patterns in unicellular eukaryotic communities (e.g., Zhang et al. 2023), the extent to which different traits related to the assembly processes of a community remains less well understood. Our results show that specific traits are differentially associated with network structure, consistent with the possibility that trait-mediated filtering is reflected in patterns of taxon association. However, these associations do not by themselves demonstrate that the traits causally generate the observed network structure but support that variation in organismal traits contributes to differences in how co-occurring taxa are organized, while network architecture likely emerges from the combined effects of multiple traits and ecological processes.

## Data availability statement

The data and code supporting the findings of this study are deposited in Zenodo. They are accessible to reviewers during the peer-review process and will be made publicly available upon publication of the manuscript.

## Acknowledgments

We thank Simone Moretti and Jonathan Jung for the precious discussions about the study. TC acknowledges funding received from the Swiss National Science Foundation (SNSF; grant 200020_201106).

## References

Adl, S.M., Bass, D., Lane, C.E., Lukeš, J., Schoch, C.L., Smirnov, A., et al. (2019). Revisions to the Classification, Nomenclature, and Diversity of Eukaryotes. Journal of Eukaryotic Microbiology, 66, 4–119.

Alve, E. (1999). Colonization of new habitats by benthic foraminifera: a review. Earth. Sci. Rev., 46, 167–185.

Balmford, A., Green, M.J.B. & Murray, M.G. (1996). Using higher-taxon richness as a surrogate for species richness: I. Regional tests. Proc. R. Soc. Lond. B Biol. Sci., 263, 1267–1274.

de Bello, F., Vandewalle, M., Reitalu, T., Lepš, J., Prentice, H.C., Lavorel, S., et al. (2013). Evidence for scale- and disturbance-dependent trait assembly patterns in dry semi-natural grasslands. Journal of Ecology, 101, 1237–1244.

Bevilacqua, S., Terlizzi, A., Claudet, J., Fraschetti, S. & Boero, F. (2012). Taxonomic relatedness does not matter for species surrogacy in the assessment of community responses to environmental drivers. Journal of Applied Ecology, 49, 357–366.

Blanchet, F.G., Cazelles, K. & Gravel, D. (2020). Co-occurrence is not evidence of ecological interactions. Ecol. Lett., 23, 1050–1063.

Bowser, S.S. (2002). Reticulopodia: Structural And Behavioral Basis For The Suprageneric Placement Of Granuloreticulosan Protists. The Journal of Foraminiferal Research, 32, 440–447.

Burkett, A.M., Huber, B.T., Young, J.R., Katz, M.E., Borrelli, C., Fung, M.K., et al. (2024). Bforams@Mikrotax: An Online Database For Deep-Sea Benthic Foraminiferal Taxonomy. J. Foraminifer. Res., 54, 394–403.

Buzas, M.A., Hayek, L.-A.C., Culver, S.J., Murray, J.W. & Jones, D.O.B. (2023). Oceanic-Scale Species Diversity of Living Benthic Foraminifera: Insights into Neogene Diversity, Community Structure, Species Duration, and Biogeography. J. Foraminifer. Res., 53, 256–265.

Buzas, M.A., Hayek, L.-A.C., Jett, J.A. & Reed, S.A. (2015). Pulsating Patches History and Analyses of Spatial, Seasonal, and Yearly Distribution of Living Benthic Foraminifera. Smithsonian Contributions to Paleobiology, iv–91.

Buzas-Stephens, P., Marchitto, T.M., Culver, S.J. & Buzas, M.A. (2025). Macroevolution In Benthic Foraminifera: A Review of Attributes Leading To Longevity. J. Foraminifer. Res., 55, 345–355.

Corliss, B. (1985). Microhabitats of benthic foraminifera within deep-sea sediments. Nature, 314, 435– 438.

Courtial, J., Lothier, J., Cukier, C., Limami, A.M. & Geslin, E. (2025). Metabolic Adaptations of Benthic Forams: Foraminiferal Species Adaptations to Intertidal Mudflat Assessed by a Metabolic Approach. Journal of Eukaryotic Microbiology, 72.

Csárdi, G., Nepusz, T., Traag, V., Horvát, S., Zanini, F., Noom, D., et al. (2006). igraph: Network Analysis and Visualization. CRAN: Contributed Packages.

Davison, J., Moora, M., Jairus, T., Vasar, M., Öpik, M. & Zobel, M. (2016). Hierarchical assembly rules in arbuscular mycorrhizal (AM) fungal communities. Soil Biol. Biochem., 97, 63–70.

Diz, P., González-Guitián, V., González-Villanueva, R., Ovejero, A. & Hernández-Almeida, I. (2023). BENFEP: a quantitative database of benthic foraminifera from surface sediments of the eastern Pacific. *Earth Syst*. Sci. Data, 15, 697–722.

Diz, P., González-Villanueva, R. & García-Roselló, E. (2024). Diversity and endemism of hard-shelled benthic foraminifera in permanently oxygen-depleted bottom waters: An analysis from the eastern Pacific. Prog. Oceanogr., 225, 103277.

Dray, S. (2016). adespatial: Multivariate Multiscale Spatial Analysis. CRAN: Contributed Packages.

Dray, S., Legendre, P. & Peres-Neto, P.R. (2006). Spatial modelling: a comprehensive framework for principal coordinate analysis of neighbour matrices (PCNM). Ecol. Modell., 196, 483–493.

Fouet, M.P.A., Schweizer, M., Singer, D., Richirt, J., Quinchard, S. & Jorissen, F.J. (2024). Unravelling the distribution of three Ammonia species (Foraminifera, Rhizaria) in French Atlantic Coast estuaries using morphological and metabarcoding approaches. Mar. Micropaleontol., 188, 102353.

Galai, G., Arbel, D., Klass, K., Grinshpan, I., Mizrahi, I. & Pilosof, S. (2024). Regional processes shape the structure of rumen microbial co-occurrence networks. Ecography, 2024.

Goetz, E.J., Greco, M., Rappaport, H.B., Weiner, A.K.M., Walker, L.M., Bowser, S., et al. (2022). Foraminifera as a model of the extensive variability in genome dynamics among eukaryotes. BioEssays, 44.

Goldstein, S. & Alve, E. (2011). Experimental assembly of foraminiferal communities from coastal propagule banks. Mar. Ecol. Prog. Ser., 437, 1–11.

Gooday, A.J. (1993). Deep-sea benthic foraminiferal species which exploit phytodetritus: Characteristic features and controls on distribution. Mar. Micropaleontol., 22, 187–205.

Gooday, A.J. (2003). Benthic Foraminifera (Protista) as Tools in Deep-water Palaeoceanography: Environmental Influences on Fauna Characteristics. Advance in Marine Biology, 46, 1–90.

Gooday, A.J., Cedhagen, T., Kamenskaya, O.E. & Cornelius, N. (2007). The biodiversity and biogeography of komokiaceans and other enigmatic foraminiferan-like protists in the deep Southern Ocean. Deep Sea Research Part II: Topical Studies in Oceanography, 54, 1691–1719.

Gooday, A.J. & Jorissen, F.J. (2012). Benthic foraminiferal biogeography: Controls on global distribution patterns in deep-water settings. Ann. Rev. Mar. Sci., 4, 237–262.

Grilli, J., Rogers, T. & Allesina, S. (2016). Modularity and stability in ecological communities. Nat. Commun., 7, 12031.

Guénard, G. & Legendre, P. (2024). Spatially explicit predictions using spatial eigenvector maps. Methods Ecol. Evol., 15, 2129–2140.

Guy-Haim, T. & Bouchet, V.M.P. (2025). Beyond taxonomy: A framework for biological trait analysis to assess the functional structure of benthic foraminiferal communities. Mar. Pollut. Bull., 213, 117699.

Henry, G. & Basso, D. (2026). Calcareous Benthic Foraminifera Under Ocean Alkalinity Enhancement: Synthesis, Knowledge Gaps, and Future Research Agenda. Earths Future, 14.

Holbourn, A., Henderson, A.S. & MacLeod, N. (2013). Atlas of Benthic Foraminifera. Wiley.

Holt, R.D. (2009). Bringing the Hutchinsonian niche into the 21st century: Ecological and evolutionary perspectives. Proceedings of the National Academy of Sciences, 106, 19659–19665.

Hoober, L., Herut, B., Teutsch, N., Ashckenazi-Polivoda, S., Torfstein, A. & Abramovich, S. (2026). Biogeochemical dichotomy and intra-order variability in Miliolid and Rotaliid foraminifera. Biogeosciences, 23, 5685–5696.

Jones, D.O.B. & Murray, J.W. (2017). Controls on the standing crop of benthic foraminifera at the oceanic scale. Mar. Ecol. Prog. Ser., 581, 71–83.

Jorissen, F.J., Meyers, S.R., Kelly-Gerreyn, B.A., Huchet, L., Mouret, A. & Anschutz, P. (2022). The 4GFOR model – Coupling 4G early diagenesis and benthic foraminiferal ecology. Mar. Micropaleontol., 170, 102078.

Jorissen, F.J., de Stigter, H.C. & Widmark, J.G.V. (1995). A conceptual model explaining benthic foraminiferal microhabitats. Mar. Micropaleontol., 26, 3–15.

Keating-Bitonti, C.R. & Payne, J.L. (2017). Ecophenotypic responses of benthic foraminifera to oxygen availability along an oxygen gradient in the California Borderland. Marine Ecology, 38.

Krivitsky, P.N., Hunter, D.R., Morris, M. & Klumb, C. (2023). ergm 4: New Features for Analyzing Exponential-Family Random Graph Models. J. Stat. Softw., 105.

Lechêne, A., Lobry, J., Boët, P. & Laffaille, P. (2018). Change in fish functional diversity and assembly rules in the course of tidal marsh restoration. PLoS One, 13, e0209025.

Leibold, M.A., Holyoak, M., Mouquet, N., Amarasekare, P., Chase, J.M., Hoopes, M.F., et al. (2004). The metacommunity concept: a framework for multi-scale community ecology. Ecol. Lett., 7, 601– 613.

Li, T., Li, B., Lin, Z., Xie, W. & Yang, C. (2025). Drift Drives Foraminiferal Community Assembly on a Carbonate Platform. Ecol. Evol., 15.

Li, T., Zhang, M., Li, B., Cai, G., Li, S. & Nie, X. (2022a). Co-occurrence patterns and community assembly mechanisms of benthic foraminiferal communities in South Chinese bays. Ecol. Indic., 144, 109489.

Li, Y., Liu, C., Sack, L., Xu, L., Li, M., Zhang, J., et al. (2022b). Leaf trait network architecture shifts with species-richness and climate across forests at continental scale. Ecol. Lett., 25, 1442–1457.

Liu, J., Xu, Z. & Liu, X. (2023). Applicability of taxonomic sufficiency of macrofaunal assemblages in benthic ecological quality assessments: Insights from a semi-enclosed bay. Front. Mar. Sci., 10.

Loeblich, A.R. & Tappan, H. (1988). *Foraminiferal Genera and Their Classification*. Springer US, Boston, MA.

Logue, J.B., Mouquet, N., Peter, H. & Hillebrand, H. (2011). Empirical approaches to metacommunities: a review and comparison with theory. Trends Ecol. Evol., 26, 482–491.

Luzuriaga, A.L., Sánchez, A.M., Maestre, F.T. & Escudero, A. (2012). Assemblage of a Semi-Arid Annual Plant Community: Abiotic and Biotic Filters Act Hierarchically. PLoS One, 7, e41270.

Mayfield, M.M. & Levine, J.M. (2010). Opposing effects of competitive exclusion on the phylogenetic structure of communities. Ecol. Lett., 13, 1085–1093.

McGill, B.J., Enquist, B.J., Weiher, E. & Westoby, M. (2006). Rebuilding community ecology from functional traits. Trends Ecol Evol, 21, 178–185.

Mistri, M. & Rossi, R. (2001). Taxonomic sufficiency in lagoonal ecosystems. Journal of the Marine Biological Association of the United Kingdom, 81, 339–340.

Moodley, L., Boschker, H., Middelburg, J., Pel, R., Herman, P., de Deckere, E., et al. (2000). Ecological significance of benthic foraminifera: 13C labelling experiments. Mar. Ecol. Prog. Ser., 202, 289– 295.

Murray, J.W. (2006). Ecology and Applications of Benthic Foraminifera. Cambridge University Press.

Murray, J.W., Alve, E. & Jones, B.W. (2011). A new look at modern agglutinated benthic foraminiferal morphogroups: their value in palaeoecological interpretation. Palaeogeogr. Palaeoclimatol. Palaeoecol., 309, 229–241.

Neff, F., Hagge, J., Achury, R., Ambarlı, D., Ammer, C., Schall, P., et al. (2022). Hierarchical trait filtering at different spatial scales determines beetle assemblages in deadwood. Funct. Ecol., 36, 2929–2942.

Nguyen, N., Pawłowska, J., Szymańska, N., Zajaczkowski, M., Weiner, A.K.M., De Schepper, S., et al. (2026). Assessing the passive dispersal of benthic foraminifera through environmental Dna. Limnol. Oceanogr., 71.

Nomaki, H., Ogawa, N.O., Ohkouchi, N., Suga, H., Toyofuku, T., Shimanaga, M., et al. (2008). Benthic foraminifera as trophic links between phytodetritus and benthic metazoans: Carbon and nitrogen isotopic evidence. Mar. Ecol. Prog. Ser., 357, 153–164.

Oksanen, J., Simpson, G.L., Blanchet, F.G., Kindt, R., Legendre, P., Minchin, P.R., et al. (2024). vegan: Community Ecology Package. CRAN: Contributed Packages.

Pawlowski, J., Fahrni, J., Lecroq, B., Longet, D., Cornelius, N., Excoffier, L., et al. (2007). Bipolar gene flow in deep-sea benthic foraminifera. Mol. Ecol., 16, 4089–4096.

Pawlowski, J., Holzmann, M., Berney, C., Fahrni, J., Gooday, A.J., Cedhagen, T., et al. (2003). The evolution of early Foraminifera. Proceedings of the National Academy of Sciences, 100, 11494– 11498.

Pawlowski, J., Holzmann, M. & Tyszka, J. (2013). New supraordinal classification of Foraminifera: Molecules meet morphology. Mar. Micropaleontol., 100, 1–10.

Ponisio, L.C., Valdovinos, F.S., Allhoff, K.T., Gaiarsa, M.P., Barner, A., Guimarães, P.R., et al. (2019). A Network Perspective for Community Assembly. Front. Ecol. Evol., 7.

Prazeres, M., Uthicke, S. & Pandolfi, J.M. (2015). Ocean acidification induces biochemical and morphological changes in the calcification process of large benthic foraminifera. Proceedings of the Royal Society B: Biological Sciences, 282, 20142782.

R Core Team. (2024). R: A Language and Environment for Statistical Computing.

Roberts, A., Austin, W., Evans, K., Bird, C., Schweizer, M. & Darling, K. (2016). A New Integrated Approach to Taxonomy: The Fusion of Molecular and Morphological Systematics with Type Material in Benthic Foraminifera. PLoS One, 11, e0158754.

Sabbatini, A., Caridi, F., Potena, D. & Negri, A. (2025). Machine Learning for identification and classification of Foraminifera: Testing on monothalamids. Mar. Micropaleontol., 195, 102442.

Saito, V.S., Cianciaruso, M.V., Siqueira, T., Fonseca-Gessner, A.A. & Pavoine, S. (2016). Phylogenies and traits provide distinct insights about the historical and contemporary assembly of aquatic insect communities. Ecol. Evol., 6, 2925–2937.

Schmiedl, G., Milker, Y. & Mackensen, A. (2023). Climate forcing of regional deep-sea biodiversity documented by benthic foraminifera. Earth. Sci. Rev., 244, 104540.

Schweizer, M., Fontaine, D. & Pawlowski, J. (2011). Phylogenetic position of two Patagonian Cibicididae (Rotaliida, Foraminifera): Cibicidoides dispars (d’Orbigny, 1839) and Cibicidoides variabilis (d’Orbigny, 1826). Revue de Micropaléontologie, 54, 175–182.

Silk, M.J. & Fisher, D.N. (2017). Understanding animal social structure: exponential random graph models in animal behaviour research. Anim. Behav., 132, 137–146.

Song, X., Gooday, A.J., Gordon, D.P., Leduc, D., Sun, Y., Wang, Z., et al. (2026). Protist-dominated hard substrate faunas thrive at the deepest ocean depths. Science (1979)., 392, 749–754.

Spasojevic, M.J. & Suding, K.N. (2012). Inferring community assembly mechanisms from functional diversity patterns: the importance of multiple assembly processes. Journal of Ecology, 100, 652– 661.

Tylianakis, J.M. & Morris, R.J. (2017). Ecological Networks Across Environmental Gradients. Annu. Rev. Ecol. Evol. Syst., 48, 25–48.

Vellend, M. (2010). Conceptual Synthesis in Community Ecology. Q. Rev. Biol., 85, 183–206.

Włodarska-Kowalczuk, M., Pawłowska, J. & Zajaczkowski, M. (2013). Do foraminifera mirror diversity and distribution patterns of macrobenthic fauna in an Arctic glacial fjord? Mar. Micropaleontol., 103, 30–39.

Zhang, H., Yang, Y., Liu, X., Huang, T., Ma, B., Li, N., et al. (2023). Novel insights in seasonal dynamics and co-existence patterns of phytoplankton and micro-eukaryotes in drinking water reservoir, Northwest China: DNA data and ecological model. Science of The Total Environment, 857, 159160.

Van Der Zwaan, G.J., Duijnstee, I.A.P., Den Dulk, M., Ernst, S.R., Jannink, N.T. & Kouwenhoven, T.J. (1999). Benthic foraminifers: proxies or problems? A review of paleocological concepts. Earth. Sci. Rev.

