## Supplementary material for "Complementary roles among traits in community assembly processes of benthic foraminifera"

**Database construction and proportion of traits**

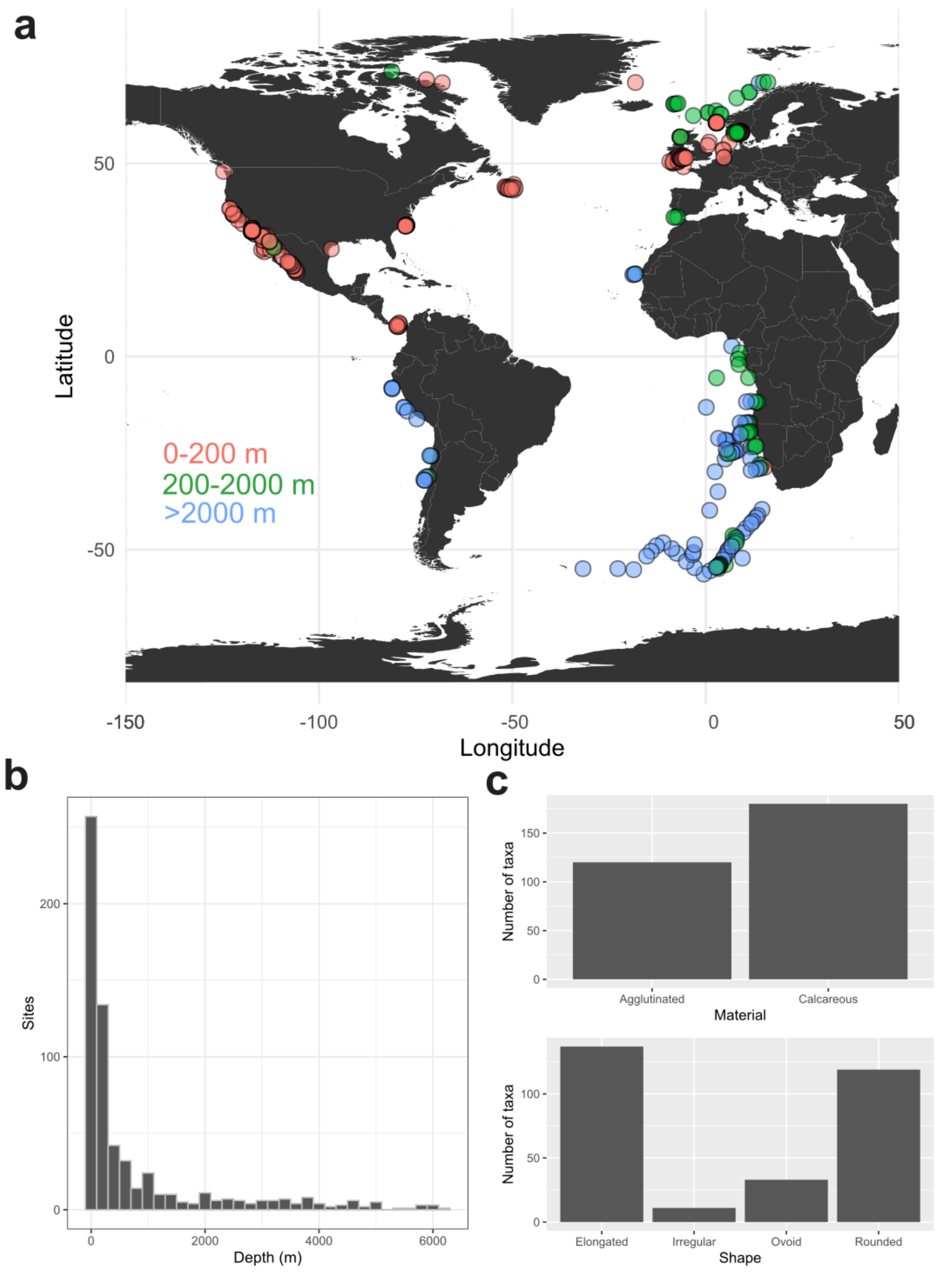


**Figure S1. General characteristics of the dataset used in this study.** a) Sample distribution, with sites colored by depth class (0-200 m = orange, 200 - 2000 = green, > 2000 m = blue). Samples were given a transparent threshold (30%) to highlight areas which were highly sampled. b) Histogram of the sites distributed along depth (bin width = 200 m). c) Number of taxa found belonging to different traits.

The database used in this study was compiled using a subset of the BENFEP_v2 (Diz et al., 2023) and the Murray (2006) datasets, by filtering the original ones for a threshold count of at least 100 living specimens. Being one of the first and most complete databases for benthic foraminifera ever done, the Murray (2006) database had 153 samples not georeferenced, while others were missing the measured depth (13). For further analyses, we remove the sites that did not have any indication of the depth of collection, along with geographical coordinates.

The Murray (2006) dataset also had some samples whose total sum of the relative abundance of the species did not sum up 100%. This is because some of the old material found in Murray (2006) have been reported in an inconsistent manner (either without decimals, or with one or two decimals, for example). To not remove most of the samples due to this artifact and to still retain most of the samples that have a good cumulative abundance, we retain only the sample with a cumulative abundance between 90 and 105 (Fig. S2).

The resulting database comprises a total of 628 sites and 297 genera. All the samples in both databases were investigated using the Rose-Bengal-stained method to detect the living component of the benthic foraminiferal fauna and are restricted only to the fractions > 63 µm. The distribution of the samples is confined to the Atlantic Ocean and the eastern Pacific coast, from 73.75° N to -56.32°N (Fig. S1a). The water depth ranges from 0.3 m to 6250 m. The average depth is 664 m, while the median depth is 104 m, reflecting the coastal nature of these sites (Fig. S1b). We also report the number of taxa belonging to agglutinated and calcareous, and to different test shapes (Fig. S1c).

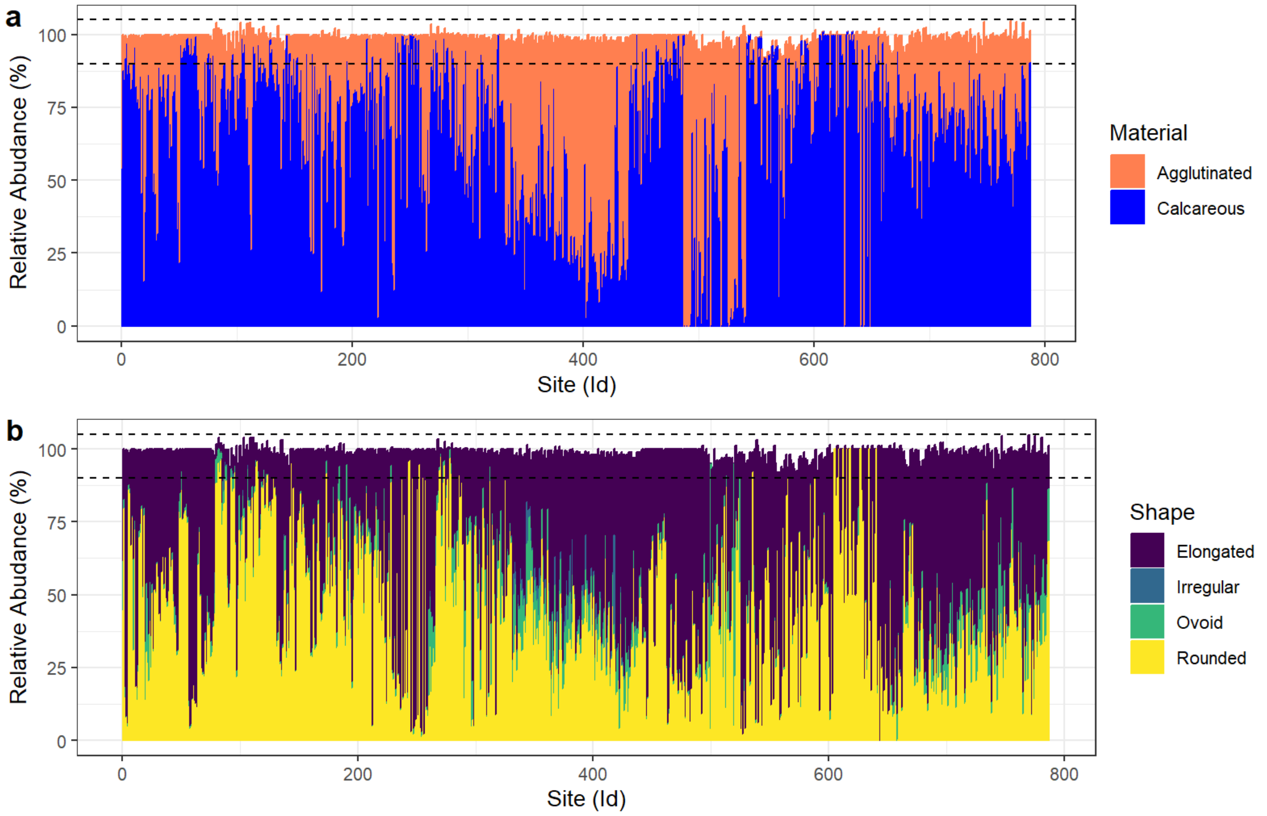


**Figure S2. Relative abundances of the two different traits categories across all the living communities analysed in this study.** a) Test-formation processes and b) General test shape.

To further assess the suitability of these datasets for inferring global-scale patterns, an incidence-based analysis (on presence/absence data) was conducted to estimate the completeness of the diversity analysed, first divided by depth bins, and then on the total datasets (*specacc* function in vegan, Fig. S3). This was done because the number of specimens per taxa was not given for every site. To maximize the information available for inference of genus co-occurrence patterns, networks were constructed using the complete set of available samples rather than by subsampling to equalize sample numbers among depth zones (Fig. S3a). Sample-based genus accumulation curves showed a pronounced decline in the rate of new genus detection with increasing sampling effort, indicating that the available dataset captured most of the observed taxonomic diversity (Fig. S3b).

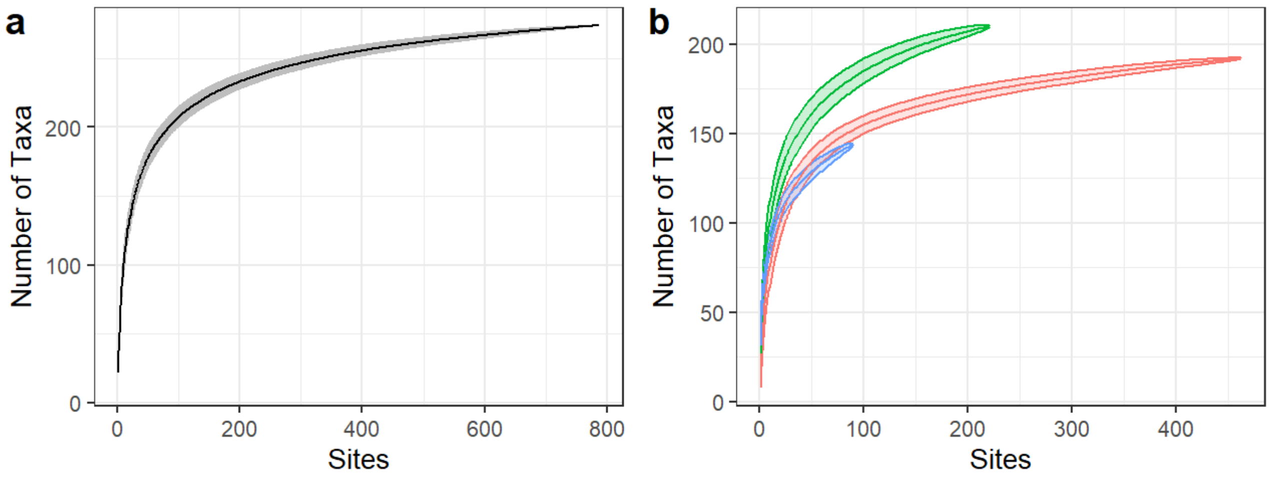


**Figure S3. Incidence-based accumulations are derived from the presence/absence community matrix.** a) Evaluated on the entirety of the datasets; b) evaluated at different depth bands (red = 0-200 m; green = 200-2000 m; blue => 2000 m).

**Spatial patterns after forward selection of the Moran Eigenvector's Map**

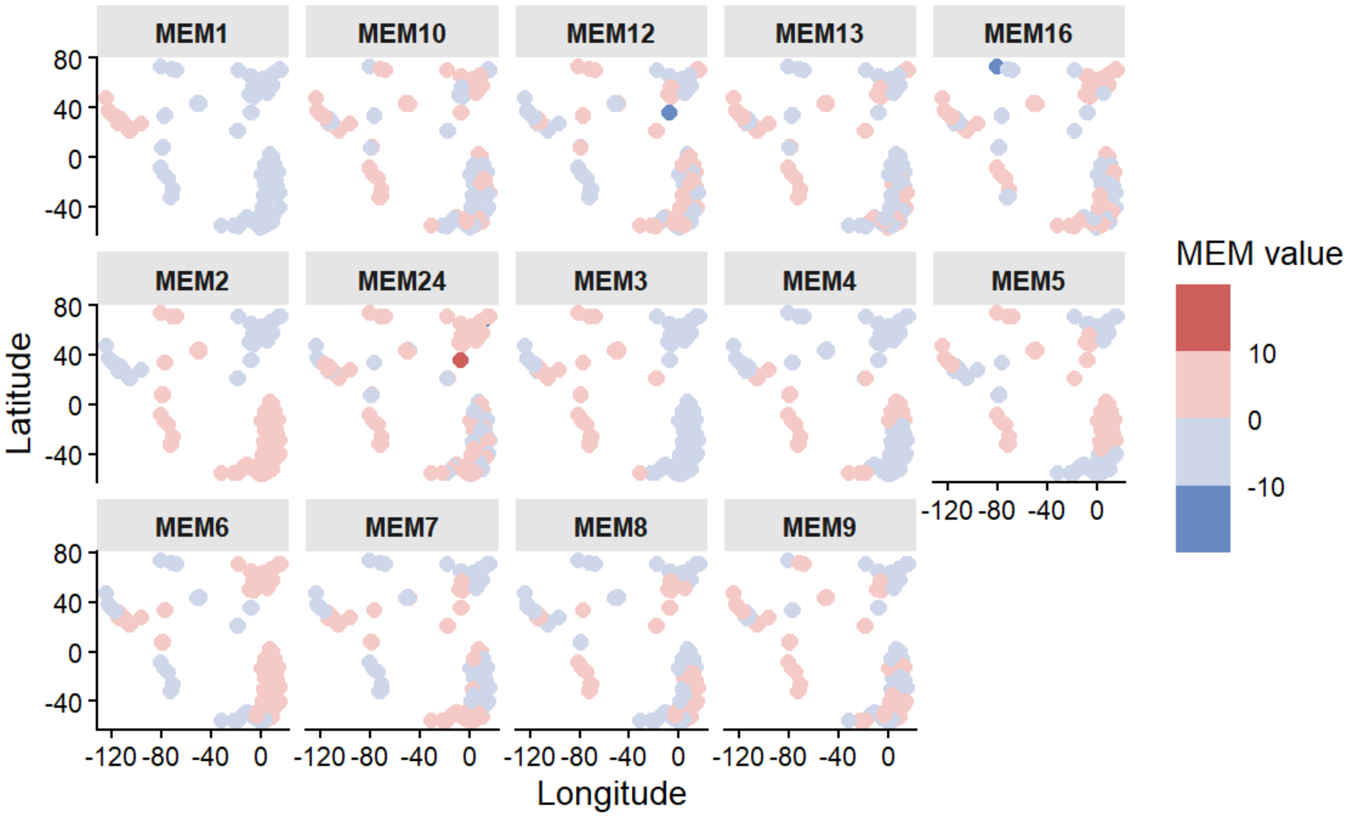


**Figure S4.** Spatial structure identified by distance-based Moran's eigenvector maps (MeMs) after forward selections. The MEM values indicate the correlations between the MeM and the spatial points. Lower MeM number corresponds to larger scale features (MeM 1 and MeM 2) while higher-order MeMs (MeM 3, MeM4 etc.) indicate the lower finer scale structure. The maps are not showing the continental margins for better visualizations.

### Constrained Redundancy Analysis

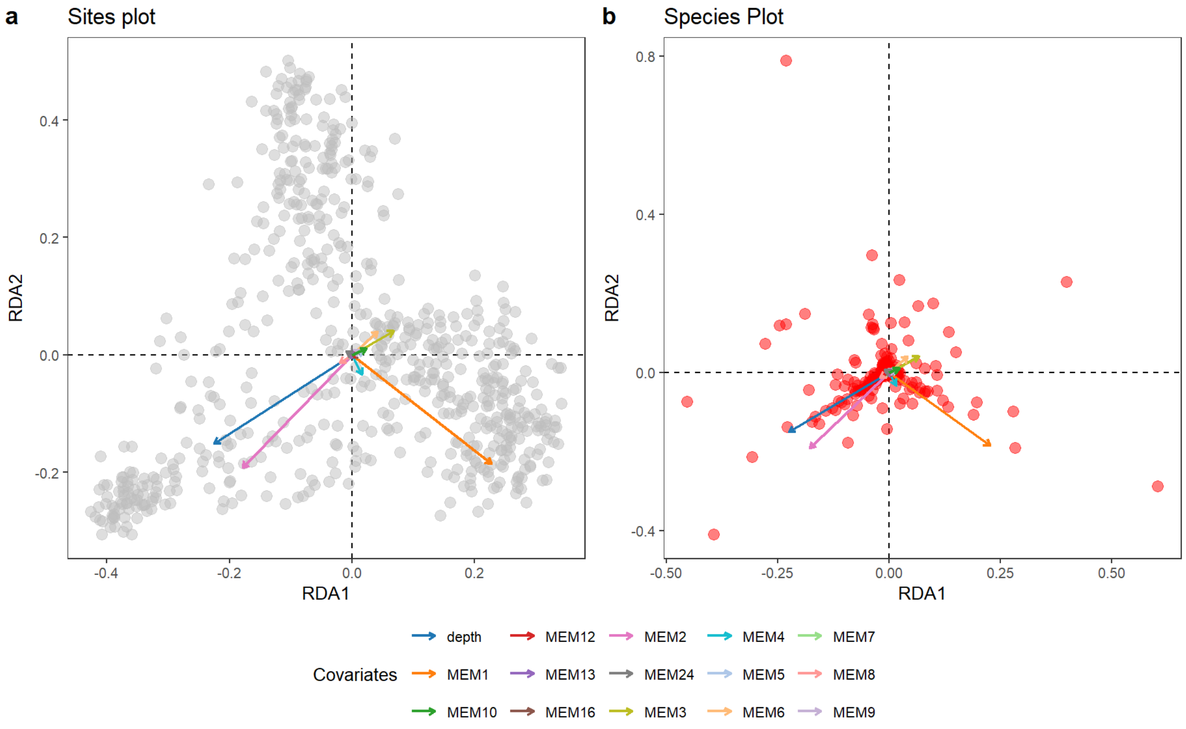


**Figure S5.** Constrained Redundancy Analysis (RDA) conducted on the communities considering geography (as MeM vectors) and Depths. For visualization, only the first 2 Moran eigenvector Maps (MeM) are shown along with the depth and all the 14 MeMs. In a) the points represent sites, while in b) they represent different taxa.

### Negative co-occurrence network

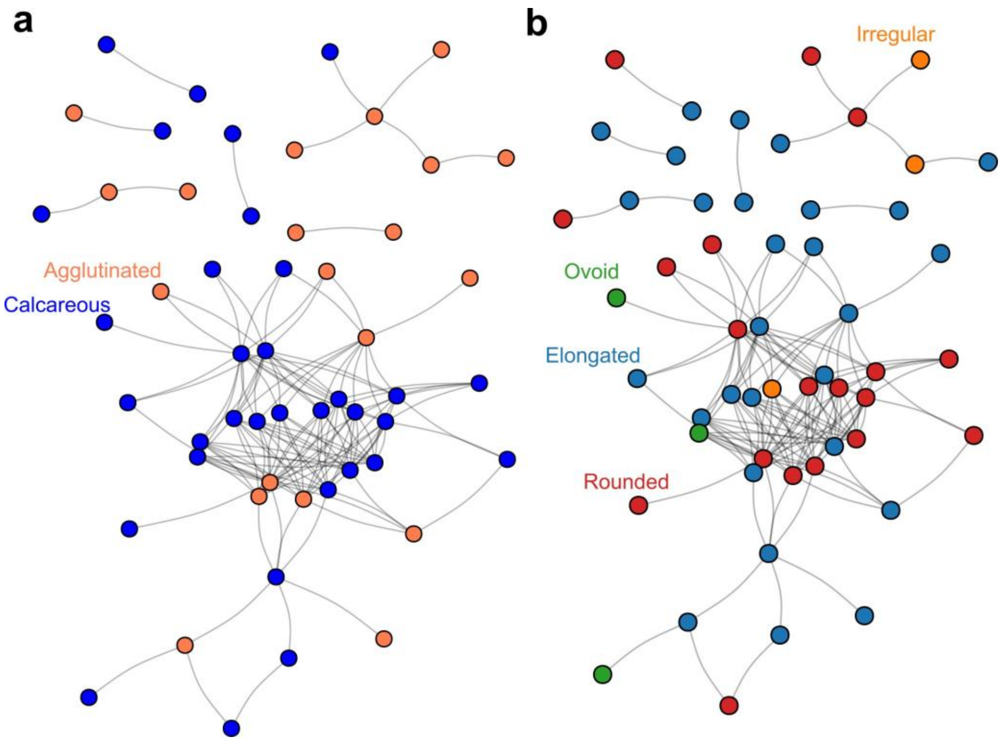


**Figure S6.** Negative co-occurrence networks derived from the correlations between the residuals extracted from the pRDA analysis. Nodes are colored as shown in Figure 2 of the main text. a) Test-construction strategies: blue = calcareous; orange = agglutinated. b) General shape: irregular = orange; ovoid = green; elongated = blue; rounded = red.

**Null-model distribution of modularity and empirical modularity**

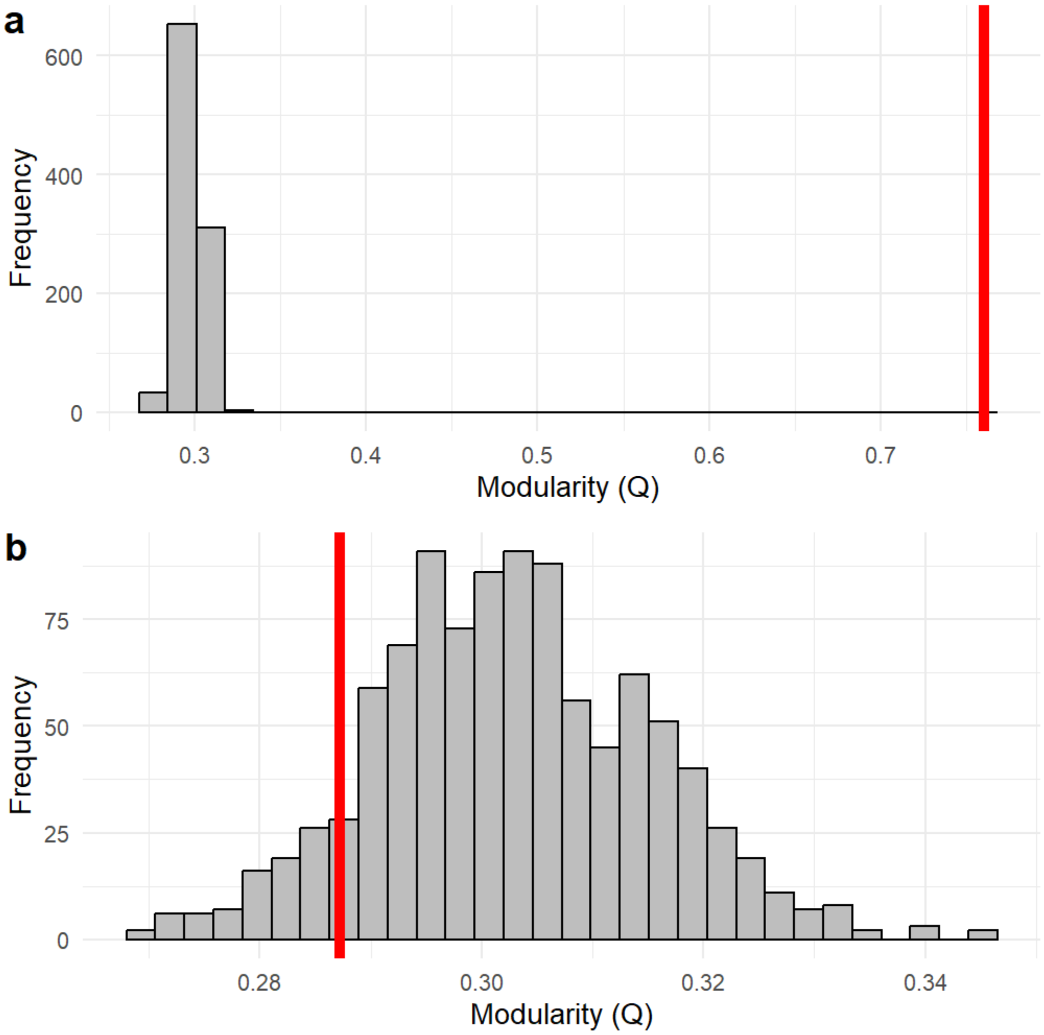


**Figure S7.** Null models result of modularity (Q) calculated on the positive (a) and negative (b) network using 999 permutations of the respective empirical networks by maintaining the number of nodes and edges. In red are shown the observed Q of the positive (a) and negative (b) one.

**Participation coefficients and degrees relationship of the positive network**

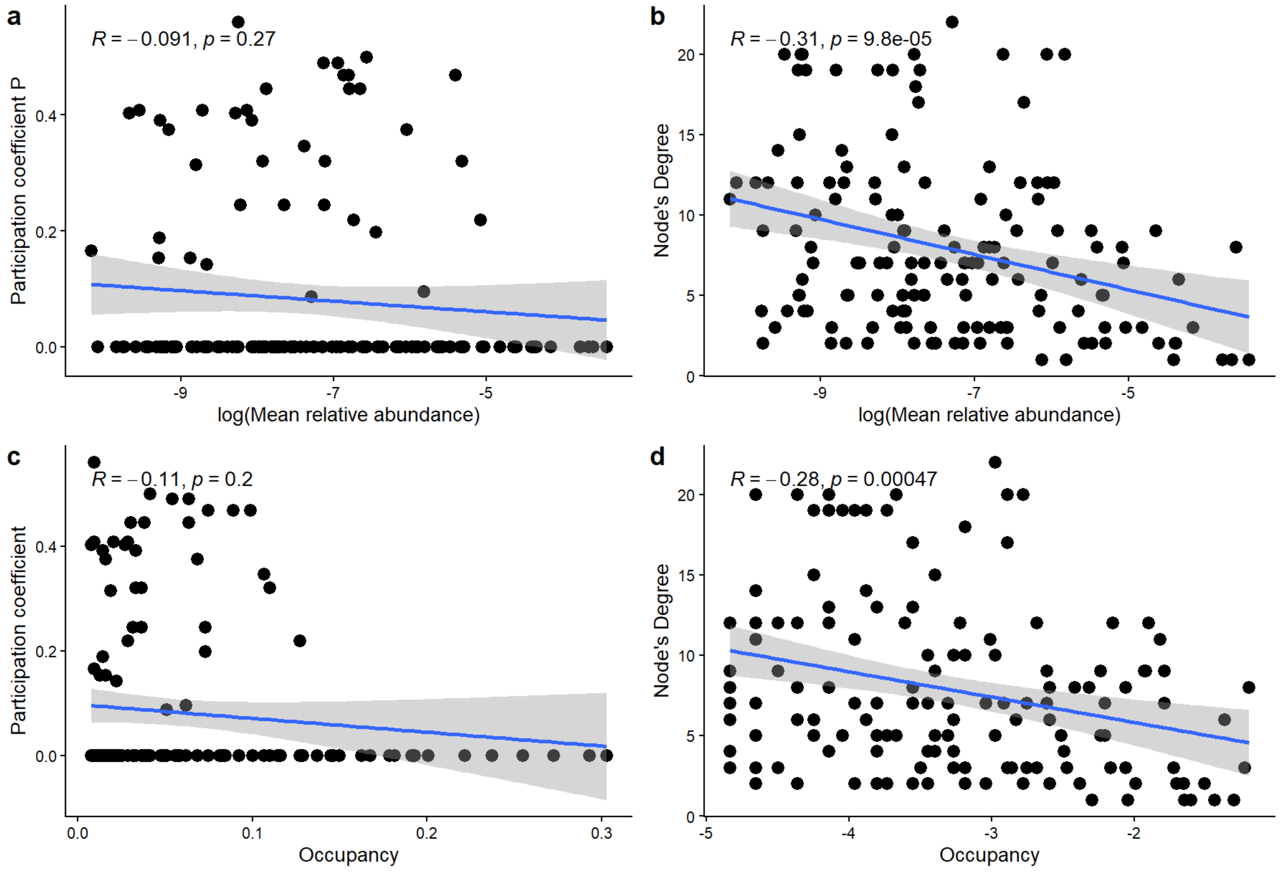


**Figure S8.** Pearson's correlations (R) between the Participation coefficient and the degree's distributions against the mean relative abundances of the taxa (a,b) and occupancy (c,d), defined as the fractional number of sites occupied. In d, the occupancy was logarithmically scaled. Participation coefficient refers to the number of edges that a node has with other modules, while the degree measures the number of edges that a node has.

**Goodness of fit of the ERGM model related to the different test-construction strategies calculated on the positive co-occurrence network.**

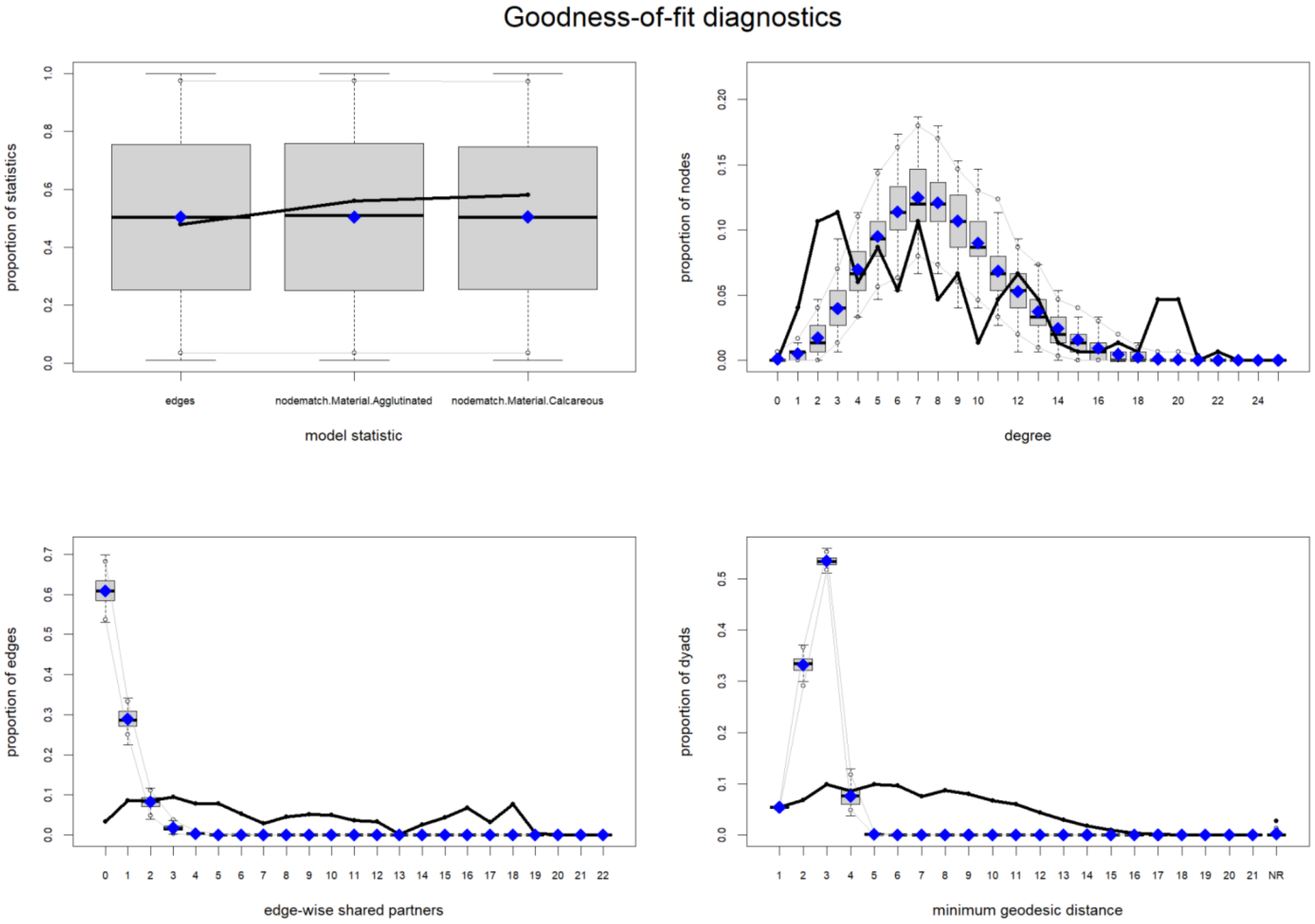


**Figure S9.** Results derived from the *gof* function applied to the ERGM referred to the positive co-occurrence network and evaluated only on the material. The empirical model is shown in black, while, in blue, is presented the aggregates of the simulations.
